# Delivery of mRNA encoding an anti-tau monoclonal antibody and engineered scFv intrabody results in functional antibody expression in SH-SY5Y cells

**DOI:** 10.1101/2023.11.15.563554

**Authors:** Patricia Wongsodirdjo, Alayna C. Caruso, Alicia Yong, Madeleine A. Lester, Laura J. Vella, Ya Hui Hung, Rebecca M. Nisbet

## Abstract

Monoclonal antibodies have emerged as a leading therapeutic agent for the treatment of disease, including Alzheimer’s disease. In the last year, two anti-amyloid monoclonal antibodies, lecanemab and aducanumab, have been approved in the USA for the treatment of Alzheimer’s disease, whilst several tau-targeting monoclonal antibodies are currently in clinical trials. Such antibodies, however, are expensive and timely to produce and require frequent dosing regimens to ensure disease-modifying effects. Synthetic *in vitro*-transcribed mRNA encoding antibodies for endogenous protein expression holds the potential to overcome many of the limitations associated with protein antibody production. Here, we have generated synthetic *in vitro*-transcribed mRNA encoding a tau specific monoclonal antibody as a full-sized IgG and as a single chain variable fragment (scFv). *In vitro* transfection of human neuroblastoma SH-SY5Y cells demonstrated the ability of the synthetic mRNA to be translated into functional tau-specific antibodies. Furthermore, we show that the translation of the tau-specific scFv as an intrabody results in the specific engagement of intracellular tau. This work highlights the utility of mRNA for the delivery of antibody therapeutics, including intrabodies, for the targeting of tau in tauopathies.

## Introduction

Alzheimer’s disease (AD) is the most common cause of dementia, a progressive neurodegenerative disease without a cure. One of the main pathological hallmarks of AD is the abnormal hyperphosphorylation and intraneuronal accumulation of the protein tau. Tau is a promising target for disease intervention and pre-clinically, tau monoclonal antibodies (mAbs) have demonstrated an ability to reduce tau pathology in several tau transgenic mouse models.^1-6^ The ability of tau mAbs to improve disease outcomes in humans, however, has not yet been demonstrated and many tau mAbs have been discontinued from clinical development.^7-10^ This is not surprising as conventional antibodies are unable to effectively cross the blood-brain barrier and enter the brain.^11,12^ Furthermore, there is limited evidence to suggest that conventional antibodies can transverse the neuronal cell membrane to engage intraneuronal tau.^13^ Conventional tau mAbs are therefore predicted to target extracellular tau and prevent tau pathology from spreading in a prion-like manner. The estimated pool of extracellular tau available for therapeutic targeting is only 0.001-0.01% of that of intracellular tau,^14,15^ however, suggesting that antibodies with enhanced targeted of intracellular tau may have better therapeutic outcomes.

Intracellular targeting of tau has been achieved in tau transgenic mouse models using intracellular antibodies (intrabodies). ^16-18^ These intrabodies were generated by engineering the single chain variable fragment (scFv) from the parental tau mAb and creating scFv-encoding adeno-associated viral vectors (AAVs). Intracerebral injection of the mice with the AAV resulted in neuronal transduction and the expressed tau intrabodies demonstrated successful reduction of pathogenic tau.^16-18^ Furthermore, comparison of an intrabody to a secreted tau scFv revealed that targeting the intracellular population of tau has an enhanced therapeutic effect.^17^ Intrabody-mediated clearance of tau has also been achieved through the addition of functional peptides that direct the complex to the proteosome or induce autophagy.^16^ Whilst viral vector-mediated intrabody gene delivery is the current preferred strategy for intrabody delivery to the brain, clinical translation of AAV use is limited due to the presence of AAV neutralizing antibodies that are generated as a result of previous exposure to wild-type virus.^19^ A promising alternative to AAV-vectored antibodies is *in vitro*-transcribed (IVT) mRNA. Synthetic IVT mRNA can be engineered at a fraction of the time and cost required to manufacture recombinant proteins and has fundamental advantages over viral-based systems, such as only requiring direct uptake into the cytosol, which results in immediate protein production. This makes mRNA a safe and fully controllable delivery tool. Such technology has been utilised pre-clinically for cancer and respiratory syncytial virus immunotherapy.^20^ Furthermore, IVT mRNA encoding single domain antibodies (V_H_H) specific for non-therapeutic targets were successfully expressed in living cells, facilitating intracellular protein targeting.^21^

Delivery of IVT mRNA encoding therapeutic antibodies and engineered intrabodies directly to neurons may transiently increase the amount of therapeutic protein in the brain compared to systemic protein delivery. We therefore wanted to determine the expression and tau engagement of our lead candidate tau antibody, RNJ1, in a conventional IgG format and as engineered scFv intrabody following IVT mRNA delivery to human neuroblastoma SH-SY5Y cells. We demonstrate that transfection of SH-SY5Y cells with mRNA encoding the light and heavy chain of RNJ1, results in expression and successful generation of secreted RNJ1 IgG, capable of binding to human tau. Furthermore, transfection of SH-SY5Y cells with mRNA encoding RNJ1 scFv, results in RNJ1 intrabody expression and engagement of intracellular tau. Together, these findings validate mRNA as an important tool for AD therapeutic development.

## Materials and Method

### Antibodies

Mouse anti-GFP antibody (Abcam) (IP: 1:200); Mouse anti-*β*-actin (Sigma-Aldrich) (WB: 1:10,000); Rabbit anti-FLAG antibody (Cell Signalling Technology) (WB: 1:1000; IF: 1:500); Mouse RNJ1 mAb (in-house) (WB: 1:1000); Tau-5 Tau-specific mAb (Millipore) (WB: 1:1000; IF: 1:500; IP: 1:1000); DAPI (Sigma-Aldrich) (IF: 1:5,000); Goat anti-mouse IRdye800CW (LI-COR) (WB: 1:10,000); Goat anti-rabbit IRdye680 (LI-COR) (WB: 1:1000).

### mRNA production

Synthetic genes (gBlocks) encoding the RNJ1 IgG heavy chain, light chain and scFv with 5’ T7 promoter and UTR sequence were generated (Integrated DNA Technologies). The scFv gene was in-frame with a C-terminal Flag tag for detection. The gBlocks were prepared following the manufacturer’s instructions and PCR amplified prior to IVT. The PCR products were purified using the QIAquick PCR Purification Kit (QIAGEN) and 0.5 *µ*g underwent IVT using the HiScribe T7 ARCA mRNA Kit with tailing (New England Biolabs) according to manufacturer’s instructions. The mRNA was purified using the Monarch RNA Cleanup Kit (NEB). A yield of approximately 30 *µ*g of mRNA was produced for each preparation.

### Cell culture

Wild-type SH-SY5Y cells and SH-SY5Y transfected with full-length human tau with a C-terminal GFP tag (Tau-GFP cells) were cultured in Dulbecco’s modified Eagle’s media (DMEM; Gibco), supplemented with 10% (v/v) fetal calf serum (FCS; Gibco) and 1% (v/v) penicillin-streptomycin (Gibco) at 37°C with 5% CO2. Tau-GFP cells were selected with 6 mg/mL blasticidin (Thermo Fisher Scientific). For expression of the RNJ1 IgG, wild-type SH-SY5Y cells were maintained in 90% (v/v) CD Hybridoma Media (Gibco), 10% (v/v) DMEM (Gibco) and supplemented with 1% (v/v) FCS and 0.1% (v/v) penicillin-streptomycin.

### mRNA transfection

Prior to transfection, SH-SY5Y cells were plated in culture media at a density of 2 x 10^5^ cells per well of a 12 well plate to achieve 80-90 % confluency at the time of transfection and incubated for 24 hrs at 37°C. Transfection was performed using Lipofectamine MessengerMax (Thermo Fisher Scientific) according to the manufacturer’s protocol. For transfection of the IgG heavy and light chain mRNA, 1.5 to 3 μg of mRNA was used. For transfection of the scFv mRNA, 0.1 to 2.5 μg of mRNA was used. To determine antibody expression, the cells and media were harvested 24 to 72 hrs post-transfection. Western blot probed with media was used neat.

### Western blotting to determine protein expression following mRNA delivery

Media collected from the SH-SY5Y cells transfected with mRNA encoding the IgG heavy chain, the IgG light chain and co-transfected with both was diluted in 4X NuPage LDS Sample Buffer (Thermo Fisher Scientific). For reduced conditions, 10X NuPage Reducing Agent (Thermo Fisher Scientific) was added and samples heated at 70°C for 10 mins. Samples were electrophoresed and western blotted with the anti-mouse secondary antibody only. To determine expression of the RNJ1 intrabody, SH-SY5Y cells transfected with the mRNA encoding the RNJ1 scFv were lysed in 1 X RIPA buffer (Abcam) and total protein concentration was measured using a BCA protein assay kit (Thermo Fisher Scientific). 3-5 *µ*g total protein was diluted in 4X NuPage LDS Sample Buffer (Thermo Fisher Scientific) with 10X NuPage Reducing Agent (Thermo Fisher Scientific) and heated at 70°C for 10 minutes. Samples were electrophoresed and western blotted with anti-Flag antibody. All membranes were imaged using the Odyssey Fc Imaging System (LI-COR). Image processing was performed on Image Studios software (LI-COR).

### Immunofluorescence

Mouse hippocampal primary neurons (DIV eight) plated out onto glass cover slips were transfected with 0.5 *µ*g mRNA encoding the RNJ1 scFv using lipofectamine2000 (Thermo Fisher Scientific). Eight days later, cells were fixed and underwent immunofluorescent labelling with anti-Flag to detect RNJ1 scFv and Tau-5 to detect tau. Cells were imaged with the Axio Observer 7 inverted microscope (Zeiss) using a 40X oil objective. System was controlled using the ZEN acquisition software. For further processing, images were z-projected by the averaging method in ImageJ processing software.

### Western blotting to determine IgG binding to Tau

Recombinant full-length human tau (hTau441) (5*μ*g) was electrophoresed and western blotted with either Tau5, RNJ1 monoclonal antibody or the undiluted media collected from SH-SY5Y cells co-transfected with mRNA encoding the RNJ1 IgG heavy chain and light chain. Membranes were imaged as described above.

### Tau pull-down assay

Tau-GFP cells were plated at a density of 6 x 10^5^ cells per well into a six-well plate in culture media and incubated overnight at 37°C. The following day, the cells were transfected with 2.5 μg of RNJ1 scFv mRNA with lipofectamine MessengerMax (Thermo Fisher Scientific) as described above. 24 hrs post transfection, cells were collected and lysed in immunoprecipitation buffer (20 mM Tris-HCl pH 7.4, 10 mM potassium chloride, 10 mM magnesium chloride, 2 mM EDTA, 10% (v/v) glycerol, 1% (v/v) Triton X-100). The lysate (“Input”) was incubated with anti-GFP antibody (1:200)(Abcam) overnight at 4°C with rotation. Following overnight incubation, 50 *µ*L protein-G agarose beads (Roche) were then added to each sample and incubated for 1 hr at room temperature with rotation. Samples were centrifuged at 5000 *x g* for 2 mins and supernatant (“flow-through”) was collected for western blotting. The pulled-down sample was washed thrice with lysis buffer and pulled down protein eluted (“IP”) by boiling in 1X NuPAGE LDS Sample Buffer (Thermo Fisher Scientific). Samples were electrophoresed and western blotted with Tau-5 antibody and anti-Flag antibody.

### Statistical analysis

Statistical analyses were performed using GraphPad Prism 9.0 software. Statistical significance between experimental groups were analysed using one-way ANOVA followed by Tukey’s multiple comparisons test, where p ≤ 0.05 was considered statistically significant. All values are reported as mean ± standard error of the mean.

## Results

### mRNA encoding RNJ1 IgG and scFv is synthesized with 5’ cap and 3’ poly(A) tail

RNJ1, our lead candidate tau-targeting mAb was used as the template for the IVT mRNA synthesis. Synthetic double-stranded DNA encoding the RNJ1 murine IgG1 heavy chain, murine kappa light chain and RNJ1 scFv fused to a triple FLAG tag, were generated. Each synthetic DNA construct contained the bacteriophage T7 promoter at the 5’ end followed by a synthetic 5’ UTR, a Kozak sequence, a transcriptional start site, and a 3’ UTR (Fig. 1A). The commonly used secretory peptide from Interleukin-2 (IL-2) was used for the secretion of the expressed RNJ1 IgG heavy and light chain ^22^. Following IVT, the mRNA contained the synthetic Anti-Reverse Cap Analog (ARCA), which prevents capping of mRNA in reverse orientation, thereby ensuring all transcripts are translatable. In addition to the 5’ ARCA cap, all mRNA constructs were synthesised to include a 3’ poly(A) tail, estimated to be 150 nts or longer (Figure 1A). Gel electrophoresis confirmed successful IVT synthesis of mRNA (Fig. 1B).

**Figure 1.**
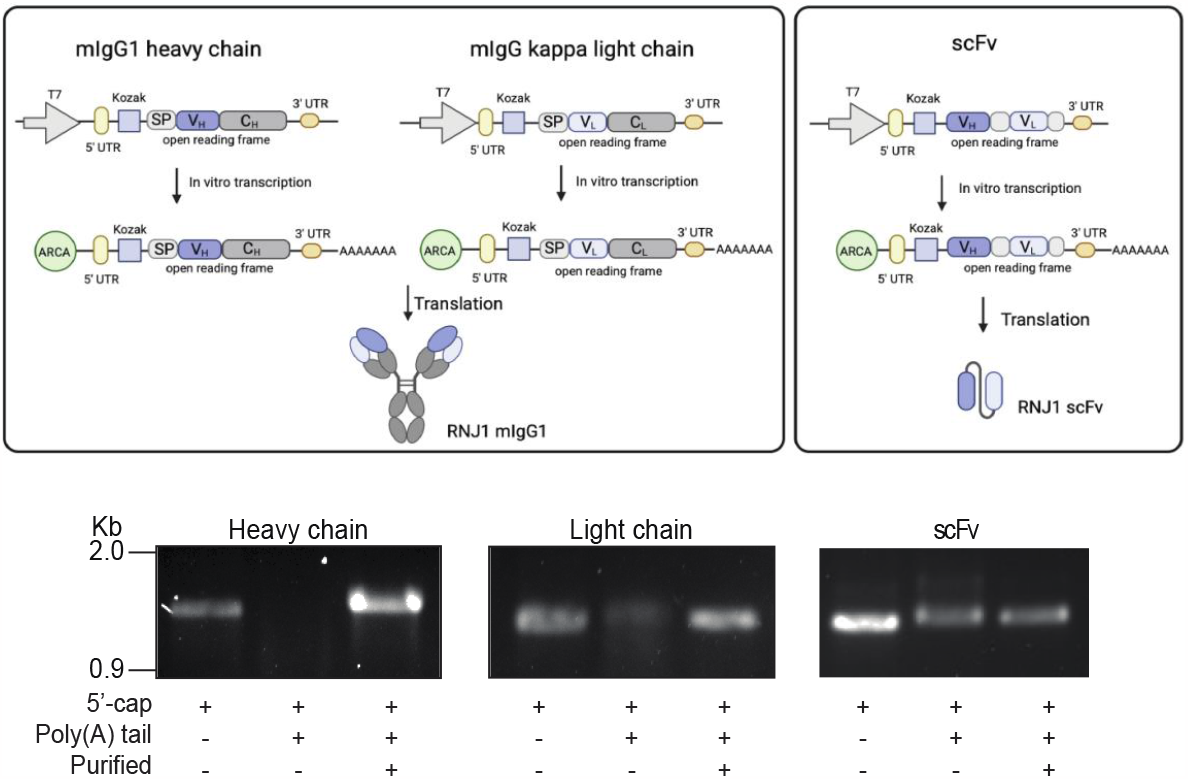
IVT synthesis of RNJ1 IgG heavy chain, light chain, and scFv mRNA. (A) Schematic of RNJ1 IgG light chain, heavy chain and scFv DNA templates and the subsequent IVT mRNA. (B) Agarose gel loaded of IVT mRNA after 5’ capping, addition of the poly(A) tail, and purification. All schematics created with Biorender.com.

### Co-transfection of IgG heavy and light chain mRNA results in the expression and secretion of functional full-length RNJ1 IgG

To determine if the delivery of mRNA encoding a mAb can result in the production and secretion of a functional antibody in human neuroblastoma cells, the mRNA encoding the mIgG heavy chain (HC) and light chain (LC) chain of RNJ1 were individually transfected or co-transfected into SH-SY5Y cells and the media collected. The RNJ1 HC and LC were both translated and secreted into the media (Fig. 2A). Western blotting revealed expression of the HC at the expected size of 50 kDa (Fig. 2A) and the LC at the expected size of 25 kDa (Fig. 2A). A HC dimer of 90 kDa was also observed (Fig. 2A). The successful generation of fulllength IgG protein via co-transfection of IVT mRNA encoding the HC and LC has previously been reported as challenging due to several methodological variables, including finding the optimal mRNA LC to HC ratio, and appropriate delivery method of the mRNA (e.g., parallel, subsequent, or simultaneous delivery).^23^ We therefore tested two different HC to LC mRNA ratios (1:1 and 1:2) using parallel co-transfection. Both conditions showed successful expression of RNJ1 at the expected size of 195 kDa (Fig. 2B). Additionally, expression levels of RNJ1 IgG followed by co-transfection at both ratios were comparable (Fig. 2B). Interestingly, an increased dose of HC and LC mRNA (3 ug of each, 1:1) showed reduced HC and LC expression and lower RNJ1 IgG levels (Fig. 2B). A reduced amount of 1.5 μg of mRNA at a 1:1 ratio of HC to LC was therefore used for subsequent transfections. Once expression of the RNJ1 mIgG1 was achieved, the ability of the mRNA encoded IgG to bind human recombinant tau was assessed. The mRNA-encoded RNJ1 mIgG was shown to detect recombinant human tau similar to the mAb version of RNJ1 IgG (Fig. 2C).

**Figure 2.**
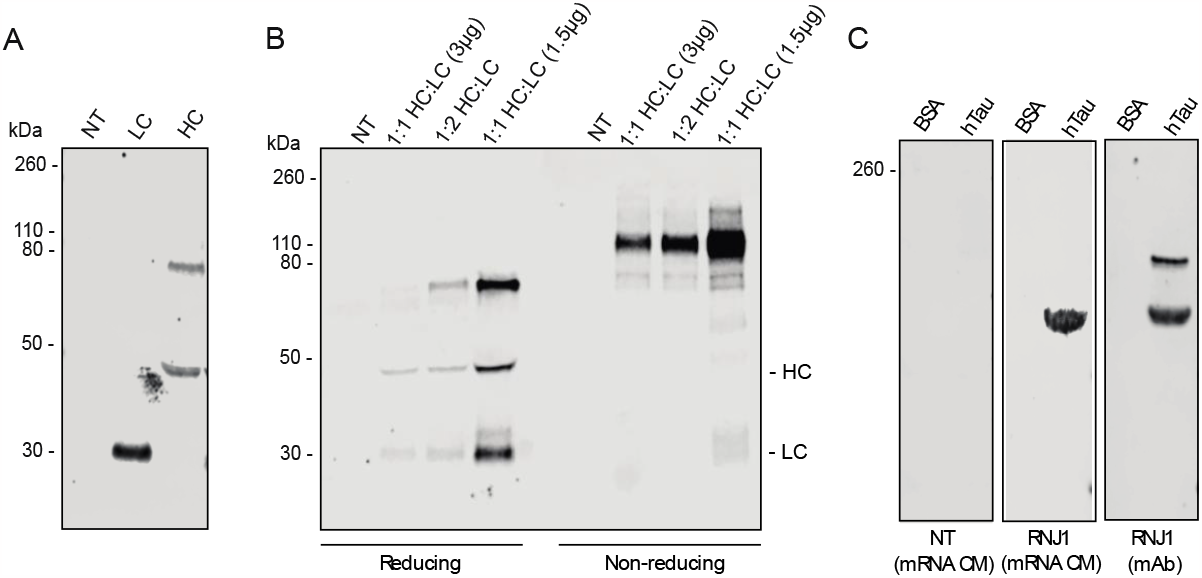
RNJ1 IgG expression and secretion following mRNA transfection. (A) Western blot of media from SH-SY5Y cells non-transfected or transfected with IgG H and/or IgG L mRNA. (B) Western blot of media collected from SH-SY5Y cells non-transfected or transfected with different ratios (1:1 or 1:2 H:L), or increased dose of IgG H to IgG L mRNA (1.5 μg or 3 μg) . (C) Western blot of recombinant human Tau probed with recombinant RNJ1 protein, or media collected from SH-SY5Y cells non-transfected or co-transfected with mRNA encoding the RNJ1 IgG H and L chain. All western blots were immunoblotted with α-rabbit secondary antibody.

### Transfection of SH-SY5Y cells with mRNA encoding RNJ1 scFv results in cytoplasmic intrabody expression and engagement of intracellular tau

Tau is predominately localised within neurons where it aggregates and induces neurotoxicity in disease. Tau antibody therapeutics may therefore have enhanced efficacy if they are able to engage intracellular tau and prevent tau aggregation. To investigate whether transfection of cells with mRNA encoding the RNJ1 scFv results in expression of a functional tau intrabody, SH-SY5Y cells were transfected with IVT mRNA encoding the RNJ1 scFv. This resulted in successful cytoplasmic generation of the RNJ1 scFv with expression levels increasing with amount of mRNA used for transfection (Fig. 3A and B). Intrabody expression was examined at 24, 48, and 72 hours post-transfection, revealing a decrease in protein expression overtime (Fig. 3C and D).

**Figure 3.**
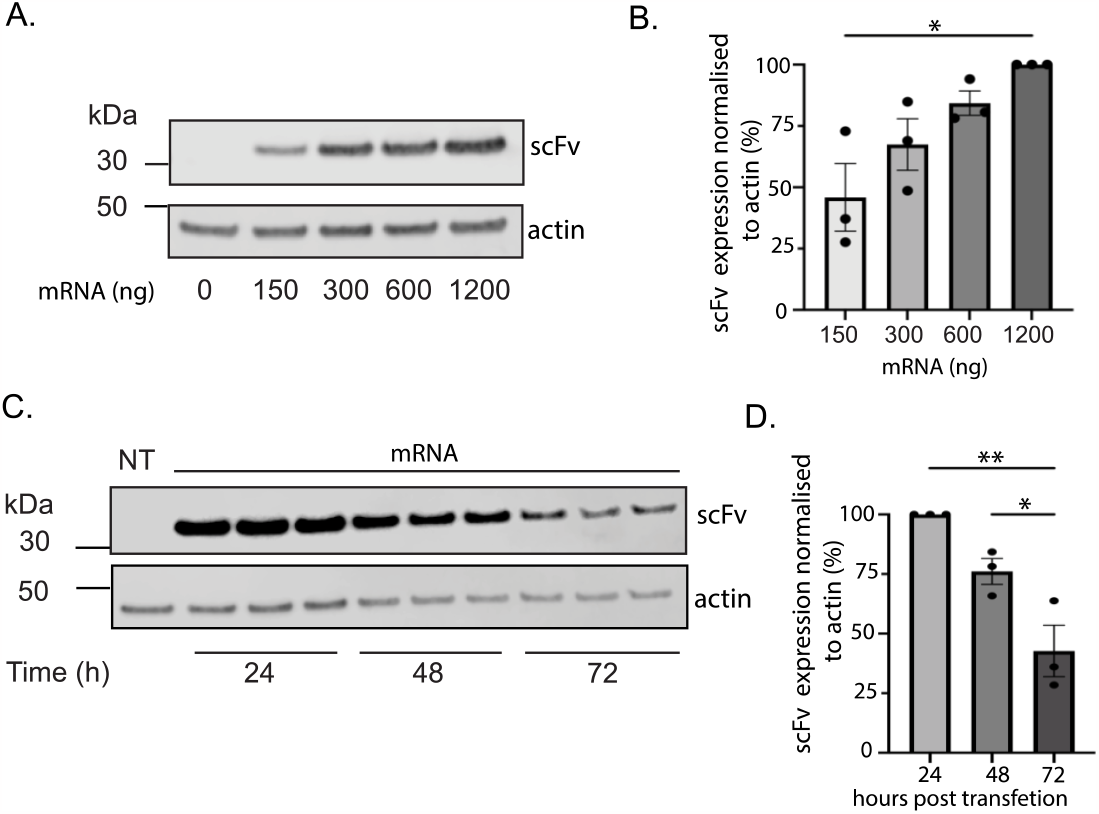
Expression of mRNA-encoded RNJ1 intrabody. (A) Representative western blot of SH-SY5Y cells non-transfected or transfected with increasing dose of RNJ1 intrabody mRNA (150-1200 ng). (B) Quantification of (A). (C) Representative western blot of SH-SY5Y cells non-transfected or transfected with RNJ1 intrabody mRNA for 24, 48, and 72 hours. (D) Quantification of (E). All western blots were immunoblotted with α-Flag antibody or β-actin where indicated. All quantification data is representative of three independent experiments (n = 3); All data points were normalised to β-actin expression; presented as mean ± SEM; analysed via one-way ANOVA with Tukey’s multiple comparisons (* = p < 0.05, ** = p < 0.01).

After expression of the RNJ1 scFv, intrabody co-localisation with and binding to intracellular tau was investigated. Following transfection of murine primary hippocampal neurons with IVT mRNA encoding the RNJ1 scFv, the RNJ1 intrabody was shown to be localized within the soma and neuronal processes of the neurons, co-localizing with endogenous murine tau (Fig. 4A). Furthermore, the interaction between the RNJ1 intrabody and cytoplasmic tau was investigated in Tau-GFP cells transfected with mRNA encoding RNJ1 scFv. The GFP-pull down assay demonstrated positive engagement between the mRNA encoded RNJ1 intrabody and intracellular tau (Fig. 4B).

**Figure 4.**
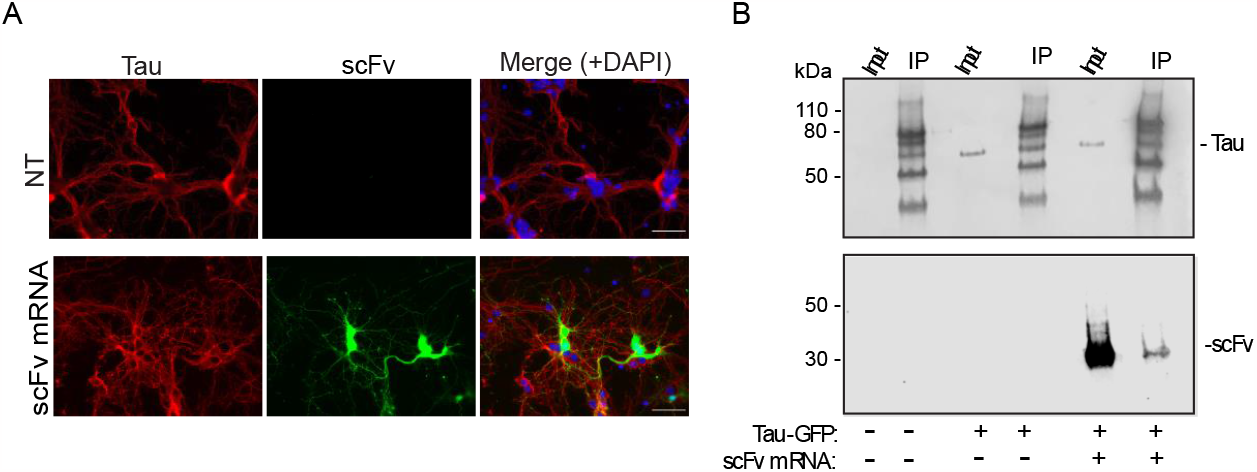
RNJ1 intrabody binds intracellular Tau. (A) Co-immunofluorescence of RNJ1 intrabody and Tau in mouse primary neurons. Neurons were probed for intracellular Tau with Tau-5 (red), RNJ1 with α-Flag (green), and DAPI (blue) as a nuclear marker. Scale bars represent 50 *µ*m. (B) Western blot of SH-SY5Y or SH-SY5Y Tau-GFP cells non-transfected or transfected with RNJ1 scFv mRNA, before (input) and after GFP-specific immunoprecipitation (IP). Immunoblot was probed with α-Flag antibody or Tau-5.

## Discussion

The clinical development of therapeutic mAbs is hindered by high costs associated with large scale production and frequent dosing regimens that are required to observe disease modifying effects. Strategies to enhance mAb bioavailability within the brain will likely improve therapeutic outcomes and reduce costs associated with the treatment of neurodegenerative diseases like AD. IVT mRNA encoding antibody therapeutics presents a promising alternative to conventional passive immunotherapy and overcomes the need to generate recombinant mAbs.

Here we demonstrate that IVT synthesised mRNA encoding a therapeutic tau antibody, RNJ1, in an IgG and scFv format, results in the endogenous translation of the antibodies when delivered to human neuroblastoma cells. Furthermore, we show that IVT mRNA encoding the RNJ1 scFv is translated in mouse primary hippocampal neurons and co-localises with tau in the cytoplasm. Importantly, the endogenously translated RNJ1 in both the full-sized IgG and smaller scFv format can engage its target, tau. For the translated RNJ1 scFv, tau engagement was shown to occur within the cell cytoplasm, making it the first documented evidence, to our knowledge, of a direct interaction between a tau antibody and tau within the cell. Whilst targeting intracellular tau, rather than extracellular tau, may be a better therapeutic strategy for the treatment of AD, full-sized IgG’s are ideal for targeting extracellular proteins such the amyloid-β peptide which accumulates to form amyloid plaques in AD. The protocol developed in this study for the effective production of IVT mRNA encoding a functional tau-targeting IgG could be applied to the development of next generation amyloid-β-specific mAbs such as lecanemab and aducanumab. From a pharmacokinetic perspective, this technology may be advantageous compared to delivery of recombinant mAbs. A recent study by Wu et al.^24^ compared the serum concentration of a bi-specific antibody targeting PD-L1 and PD-1 for the treatment of intestinal cancer, delivered as IVT mRNA or a recombinant protein and found that a single injection of the bi-specific antibody mRNA, encapsulated within lipid nanoparticles, resulted in more robust serum antibody levels with enhanced duration compared the recombinant bi-specific antibody from mammalian cell culture sources.^24^

It is important to highlight that the IVT mRNA in this study was generated from synthetic double-stranded DNA templates, rather than linearized plasmids. This strategy overcomes the need for time-consuming and expensive plasmid-based cloning which allows for a more simplified and rapid process. This would therefore allow mRNA sequence variants for enhanced translation, stability and functionality to be easily synthesised.^25^ This may include substituting the RNA bases cystidine and uracil with 5-methyl-cytidine-5′-triphosphate and pseudouridine-5′-triphosphate, respectively to further reduce immunogenicity and cytotoxicity in the host or adding the poly(A) tail to the template to ensure the addition of a poly(A) tail at a pre-determined length. Antibody secretion can also be improved using an alternative signal peptide. A study by Rybakova et al.^23^ compared multiple signal peptides to enhance serum levels of a therapeutic anti-HER2 monoclonal antibody, trastuzumab, following IVT mRNA delivery *in vivo*. ^23^ Their study revealed that although intracellular levels of trastuzumab were not affected by the signal peptide sequence, the mRNA containing human Ig kappa light chain signal peptide resulted in higher levels of trastuzumab in the cell medium compared to trastuzumab with GLuc and H5/L1 signal peptides.^23^ Furthermore, functionalization of intrabodies can be achieved by adding with protein degradation moieties to the scFv sequence for enhanced clearance of the target protein.^16,26^

Finally, *in vivo* mRNA immunotherapy studies have been limited to targeting peripheral targets^23,24,27,28^ as mRNA is predominantly encapsulated in lipid nanoparticles which are unable to effectively enter the brain. Delivery to the central nervous system would require packaging of the mRNA into vehicles capable of crossing the blood-brain barrier. To date, mRNA delivery to the brain following intravenous administration has not been achieved and therefore, *in vivo* studies have utilised intracranial injection of the mRNA into the brain.^29^ Unfortunately, this is a significant hurdle of the translation of mRNA neurotherapeutics to the clinic and future work should be directed at optimising mRNA delivery across the blood-brain barrier. In conclusion, we show that IVT synthesised mRNA can be used to generate both functional fulllength IgG antibodies and scFv intrabodies targeting tau. Our study demonstrates the utility of mRNA as a delivery platform for antibody therapeutics.

## Data availability statement

The data that support the findings of this study are available from the corresponding author, upon reasonable request.

## Acknowledgements

We thank Dr Sarah Gordon and her team for providing the mouse primary neurons and assisting with microscopy. We thank Celeste Mawal for managing the lab. We acknowledge the traditional custodians of the land on which this work was conducted, the Wurundjeri people of the Kulin nation.

## Author contributions

P.W: Methodology, investigation, formal analysis, visualization, writing. A.C.C: Methodology, investigation, visualization, and writing; A.Y: investigation, visualization and writing. M.L: investigation. L.J.V: Supervision. Y.H.H: Methodology and supervision. R.M.N: Supervision, conceptualization, methodology, formal analysis, visualization, writing and editing.

## References

1. Bajracharya R, Cruz E, Götz J, Nisbet RM. Ultrasound-mediated delivery of novel tau-specific monoclonal antibody enhances brain uptake but not therapeutic efficacy. Journal of Controlled Release. 2022/09/01/ 2022;349:634–648. 10.1016/j.jconrel.2022.07.026

2. d’Abramo C, Acker CM, Jimenez HT, Davies P. Tau passive immunotherapy in mutant P301L mice: antibody affinity versus specificity. PLoS One. 2013;8(4):e62402. doi:10.1371/journal.pone.0062402

3. Castillo-Carranza DL, Sengupta U, Guerrero-Munoz MJ, et al. Passive immunization with Tau oligomer monoclonal antibody reverses tauopathy phenotypes without affecting hyperphosphorylated neurofibrillary tangles. J Neurosci. Mar 19 2014;34(12):4260–72. doi:10.1523/JNEUROSCI.3192-13.2014

4. Umeda T, Eguchi H, Kunori Y, et al. Passive immunotherapy of tauopathy targeting pSer413-tau: a pilot study in mice. Ann Clin Transl Neurol. Mar 2015;2(3):241–55. doi:10.1002/acn3.171

5. Ittner A, Bertz J, Suh LS, Stevens CH, Gotz J, Ittner LM. Tau-targeting passive immunization modulates aspects of pathology in tau transgenic mice. J Neurochem. Jan 2015;132(1):135–45. doi:10.1111/jnc.12821

6. Boutajangout A, Ingadottir J, Davies P, Sigurdsson EM. Passive immunization targeting pathological phospho-tau protein in a mouse model reduces functional decline and clears tau aggregates from the brain. J Neurochem. Aug 2011;118(4):658–67. doi:10.1111/j.1471-4159.2011.07337.x

7. Hoglinger GU, Litvan I, Mendonca N, et al. Safety and efficacy of tilavonemab in progressive supranuclear palsy: a phase 2, randomised, placebo-controlled trial. Lancet Neurol. Mar 2021;20(3):182–192. doi:10.1016/S1474-4422(20)30489-0

8. Koga S, Dickson DW, Wszolek ZK. Neuropathology of progressive supranuclear palsy after treatment with tilavonemab. Lancet Neurol. Oct 2021;20(10):786–787. doi:10.1016/S1474-4422(21)00283-0

9. Dam T, Boxer AL, Golbe LI, et al. Safety and efficacy of anti-tau monoclonal antibody gosuranemab in progressive supranuclear palsy: a phase 2, randomized, placebo-controlled trial. Nat Med. Aug 2021;27(8):1451–1457. doi:10.1038/s41591-021-01455-x

10. Kim B, Mikytuck B, Suh E, et al. Tau immunotherapy is associated with glial responses in FTLD-tau. Acta Neuropathol. Aug 2021;142(2):243–257. doi:10.1007/s00401-021-02318-y

11. Atwal JK, Chen Y, Chiu C, et al. A therapeutic antibody targeting BACE1 inhibits amyloid-beta production in vivo. Sci Transl Med. May 25 2011;3(84):84ra43. doi:10.1126/scitranslmed.3002254

12. Levites Y, Smithson LA, Price RW, et al. Insights into the mechanisms of action of anti-Abeta antibodies in Alzheimer’s disease mouse models. FASEB J. Dec 2006;20(14):2576–8. doi:10.1096/fj.06-6463fje

13. Congdon EE, Chukwu JE, Shamir DB, et al. Tau antibody chimerization alters its charge and binding, thereby reducing its cellular uptake and efficacy. EBioMedicine. Apr 2019;42:157–173. doi:10.1016/j.ebiom.2019.03.033

14. Takeda S, Commins C, DeVos SL, et al. Seed-competent high-molecular-weight tau species accumulates in the cerebrospinal fluid of Alzheimer’s disease mouse model and human patients. Ann Neurol. Sep 2016;80(3):355–67. doi:10.1002/ana.24716

15. Bespalov A, Courade JP, Khiroug L, Terstappen GC, Wang Y. A call for better understanding of target engagement in Tau antibody development. Drug Discov Today. Nov 2022;27(11):103338. doi:10.1016/j.drudis.2022.103338

16. Gallardo G, Wong CH, Ricardez SM, et al. Targeting tauopathy with engineered tau-degrading intrabodies. Mol Neurodegener. Oct 22 2019;14(1):38. doi:10.1186/s13024-019-0340-6

17. Goodwin MS, Sinyavskaya O, Burg F, et al. Anti-tau scFvs Targeted to the Cytoplasm or Secretory Pathway Variably Modify Pathology and Neurodegenerative Phenotypes. Mol Ther. Feb 3 2021;29(2):859–872. doi:10.1016/j.ymthe.2020.10.007

18. Ising C, Gallardo G, Leyns CEG, et al. AAV-mediated expression of anti-tau scFvs decreases tau accumulation in a mouse model of tauopathy. J Exp Med. May 1 2017;214(5):1227–1238. doi:10.1084/jem.20162125

19. Rapti K, Louis-Jeune V, Kohlbrenner E, et al. Neutralizing antibodies against AAV serotypes 1, 2, 6, and 9 in sera of commonly used animal models. Mol Ther. Jan 2012;20(1):73–83. doi:10.1038/mt.2011.177

20. Tiwari PM, Vanover D, Lindsay KE, et al. Engineered mRNA-expressed antibodies prevent respiratory syncytial virus infection. Nat Commun. Oct 1 2018;9(1):3999. doi:10.1038/s41467-018-06508-3

21. Zhou X, Hao R, Chen C, et al. Rapid Delivery of Nanobodies/V(H)Hs into Living Cells via Expressing In Vitro-Transcribed mRNA. Mol Ther Methods Clin Dev. Jun 12 2020;17:401–408. doi:10.1016/j.omtm.2020.01.008

22. Zhang L, Leng Q, Mixson AJ. Alteration in the IL-2 signal peptide affects secretion of proteins in vitro and in vivo. J Gene Med. Mar 2005;7(3):354–65. doi:10.1002/jgm.677

23. Rybakova Y, Kowalski PS, Huang Y, et al. mRNA Delivery for Therapeutic Anti-HER2 Antibody Expression In Vivo. Mol Ther. Aug 7 2019;27(8):1415–1423. doi:10.1016/j.ymthe.2019.05.012

24. Wu L, Wang W, Tian J, et al. Engineered mRNA-expressed bispecific antibody prevent intestinal cancer via lipid nanoparticle delivery. Bioengineered. Dec 2021;12(2):12383–12393. doi:10.1080/21655979.2021.2003666

25. Tossberg JT, Esmond TM, Aune TM. A simplified method to produce mRNAs and functional proteins from synthetic double-stranded DNA templates. Biotechniques. Oct 2020;69(4):281–288. doi:10.2144/btn-2020-0037

26. Butler DC, Messer A. Bifunctional anti-huntingtin proteasome-directed intrabodies mediate efficient degradation of mutant huntingtin exon 1 protein fragments. PLoS One. 2011;6(12):e29199. doi:10.1371/journal.pone.0029199

27. Panova EA, Kleymenov DA, Shcheblyakov DV, et al. Single-domain antibody delivery using an mRNA platform protects against lethal doses of botulinum neurotoxin A. Front Immunol. 2023;14:1098302. doi:10.3389/fimmu.2023.1098302

28. Li JQ, Zhang ZR, Zhang HQ, et al. Intranasal delivery of replicating mRNA encoding neutralizing antibody against SARS-CoV-2 infection in mice. Signal Transduct Target Ther. Oct 25 2021;6(1):369. doi:10.1038/s41392-021-00783-1

29. Peng H, Guo X, He J, et al. Intracranial delivery of synthetic mRNA to suppress glioblastoma. Mol Ther Oncolytics. Mar 17 2022;24:160–170. doi:10.1016/j.omto.2021.12.010

